# Development of honeybee waggle dance and its differences between recruits and scouts

**DOI:** 10.1101/179408

**Authors:** Hiroyuki Ai, Yuuki Kobayashi, Toshiyuki Matake, Shinya Takahashi, Koji Hashimoto, Sakashi Maeda, Naoyuki Tsuruta

**Author notes:** Corresponding author: Hiroyuki Ai.

## Abstract

The lifetime development of the waggle dance of 14 honeybees was automatically recorded just after the imaginal molt using high-definition camera modules connected with a Raspberry Pi computer and numbered radio-frequency identification tags fitted to the back of each bee. For most honeybees, waggle dance follow preceded the appearance of the first waggle dance from 1 week after the imaginal molt. The duration per trip increased just after waggle dance follow. Before the appearance of the first waggle dance, the honeybee repeatedly follows waggle dances that indicate a limited number (2–6) of food source locations. We discriminated between two types of foragers with different roles, recruits and novice scouts, by comparing the vectors indicated by the bees’ first waggle dance (sending vectors) with dances they had previously followed (received vectors). Of 14 tagged honeybees, 11 were categorized as recruits and 2 as novice scouts. For recruits (but not for novice scouts), the duration per trip increased significantly after waggle dances follow and substantially increased just before the appearance of the first waggle dance. Moreover, recruits increased the number of times they followed waggle dances indicating the same location, and their first waggle dance indicated this location. These results suggest that the differentiation of these two types of foragers is partly related to behavioral differences after waggle dance follows: whether trip is activated or not by follows a waggle dance.

**Summary statement:** Because of technological difficulties, there are no studies comparing the development of recruit and scout waggle dances. Using miniature radio frequency identification tags, we observed and clarified these developmental processes.

**List of Abbreviations:**

AN: antenna
RFID: radio frequency identification.

## Introduction

Honeybees communicate in the hive to share foraging information and to recruit hive mates to visit profitable flowers. The waggle dance allows the communication of spatial information about food sources (von Frisch 1965; Mautz 1971; Bozic and Valentincic 1991; Judd, 1995). The effect of the waggle dance on social foraging has been evaluated theoretically and behaviorally (Seeley, 1983; Seeley and Visscher, 1988, Okada et al., 2008). The dance consists of a waggle phase, during which the dancer walks straight along the run axis and waggles the abdomen, and a return phase, during which the dancer returns to the starting position of the waggle run and usually repeats these movements (von Frisch, 1965). Measurement of flight trajectories using a harmonic radar have shown that the dance followers fly according to the vector information indicated by the waggle run (Riley et al, 2005).

The waggle dance communicates not only spatial information but also olfactory cues (von Frisch, 1967; Thom et al, 2007). Olfactory cues about food sources are communicated in the hive through trophallaxis (von Frisch, 1967), which occurs not only among foragers and receivers, but among all hive mates, resulting in propagation of the colony (Grüter et al, 2006). Learning the scent of flowers at a young age through trophallaxis may facilitate the development of a specific waggle dance; additionally, trophallaxis may produce long-term memories of profitable food sources (Balbuena et al., 2012). Thus, it has been suggested that trophallaxis experiences can affect waggle dance communications. However, the development of behaviors related to the waggle dance after the imaginal molt has not been clarified.

Most foragers receive spatial information about a flower by following the waggle dance and then visiting the flower; these are the recruits. In contrast, a small number of foragers independently visit flowers without using information from the waggle dance; these are the scouts (Oettingen-Spielberg, 1949; Lindauer, 1952). Seeley (1983) suggests that the proportion of scouts is greater in experienced foragers than in novice foragers, but this proportion also depends on foraging conditions. Seeley and Visscher (1988) suggest scouts show shorter trip durations than recruits and that scouts take smaller amounts of nectar and pollen back to the hive than recruits. However, the behavioral characteristics of recruits and scouts during the behavioral development process have not been evaluated. In this study, we investigated the development of waggle dance-related behaviors by long-term tracking of radio-frequency identification (RFID)-tagged individuals after the imaginal molt. We compared the developmental process in scouts and recruits.

## Materials and Methods

### Animals

An observation hive was built on 31 July, 2015, in the Fukuoka University campus; this contained a queen, 1000 workers, and two combs. The bees were left to settle in this observation hive (Fig. 1A and B). We began the experiments 2 months after establishing the hive when the colony size was small enough to record all behaviors visually and to clearly recognize the numbers on the backs of all honeybees in the colony. Numbers ranging from 1 to 100 were input into 100 RFID tags. The tag size was 2.5 mm × 2.5 mm × 0.4 mm and the tag weight was 5.6 mg. Then each RFID tag was sealed with a number that was the same as the RFID tag’s ID. Two combs that contained sealed cells with pupae were incubated at 33°C on 10 September 2015. On 11 September, 100 bees that emerged from the sealed cells were anesthetized and an RFID tag was fitted to the dorsal thorax of each bee using skin adhesive (SAUER-Hautkleber, Manfred Sauer GmbH, Germany, Fig. 1C). When the bees were in a state of arousal, they were put into the observation hive.

**Fig. 1:**
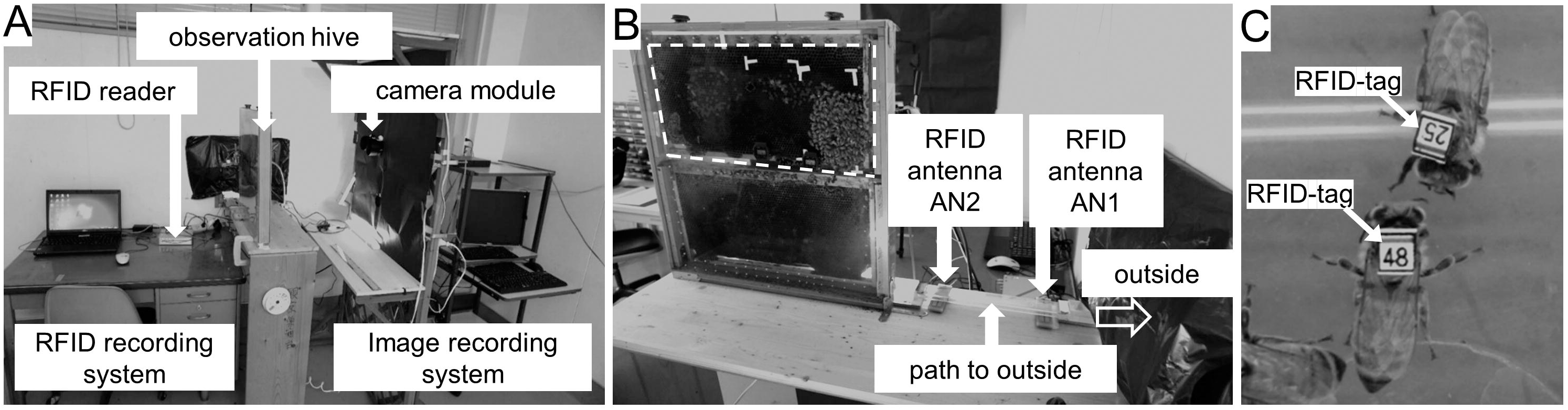
Automatic recording system for honeybee behavior. A, The system comprised an RFID recording system (on the left) and image recording system (on the right). During the recording three camera modules were located on both sides of the observation hive (in A the camera module on the left of the observation hive has been removed to show the RFID recording system). B, Behaviors of all honeybees in the colony were recorded by the camera modules. During the recording the colony was located on the upper comb (surrounded by white broken line). Two RFID antennas were located on the entrance path of the observation hive to detect the time passing through and walking directions. C, Numbered RFID-tagged honeybees. Just after emerging from the pupa, each honeybee was fitted on the back with a numbered, sealed tag.

### Behavior monitoring system

Our behavioral monitoring system consists of an RFID recording system and image recording system (Fig. 1A). Two RFID antennas (UHF RFID antenna AN-UDUL1 (920), Sobal Co. Ltd., Japan) were located on the entrance path to the observation hive (Fig. 1B). Two RFID antennas were connected with the RFID readers (UHF RFID reader, UP4-1000-J2, Sobal Co. Ltd.) and the RFID logs were stored in a personal computer. The camera modules (Raspberry Pi2 HD, Raspberry Pi Foundation, UK) were located on both sides of the observation hive. Three cameras were used to record the whole colony. The pixel count of one camera was 1296 × 730 and the acquisition frame rate was 49 fps. The covered area of one camera on the observation comb was set at 10 cm × 17 cm; this allowed visual recognition of the numbers on the bees’ backs. RFID and video recordings were obtained from 6:30 to 19:30 per day, from 14 September to 6 October 2015. The video data and RFID logs were stored in a LAN server in Fukuoka University. The RFID logs and in-hive behaviors were observed and analyzed manually using a video LAN client media player.

### RFID log analyses

On the RFID monitoring system, we could record the time the bee passed through the RFID antennas. Two RFID antennas (AN1 and AN2) were located along the entrance path to the observation hive (Fig. 1B): AN1 was set close to the outside and AN2 close to the observation hive. We could distinguish between departure and homing by the time delay in passing through these two antennas. We recorded the departure time, homing time, and duration per trip from the RFID logs of each tagged bee.

### Analyses of waggle dance-related behaviors

Using the image recording system, we recorded the time at which waggle dance-related behaviors appeared, trophallaxis, tremble dances, follows, and waggle dances. The waggle dance usually repeats several cycles of a waggle phase and a return phase and the number of cycles depends on the profitability of the flower (von Frisch, 1963). We defined movements with more than two cycles as a “waggle dance.” We discriminated between the “waggle dance” and movements with one cycle, which we termed a “one-cycle waggle dance.” In addition, we recorded the following behaviors:

1. No follow: encountered the waggle dance, then did not follow the dance.
2. One-cycle follow: encountered the waggle dance, then followed the one-cycle waggle dance.
3. Follow: encountered the waggle dance, then followed the waggle dance.

### Estimating the location of indicated flowers

Fourteen honeybees that performed their first waggle dance (Nos. 02, 05, 24, 29, 34, 46, 47, 57, 65, 70, 88, 89, 93, 99) were used to analyze the vector information for the first waggle dance and for the previous waggle dances they had followed, and to estimate the locations indicated by these waggle dances. The procedure for estimating the locations was as follows: During the waggle dance on the vertical comb, the body orientation during the waggle phase indicates the direction of the food source relative to the sun (von Frisch, 1967). In this study, the sun direction at the time at which the waggle dances occurred was obtained from the following calculation website: http://keisan.casio.jp/has10/SpecExec.cgi?id=system/2006/1185781259

The time during the waggle phase changes linearly with the distance to the location indicated by the waggle dancer (y = 0.7946x + 0.4121, duration of waggle phase (y), distance to the location (x), calculated from the data in von Frisch, 1967). The distances were calculated using this equation. The points obtained from vectors indicated by the waggle dance were drawn in the polar coordinates around the observation hive.

To evaluate whether the points were encoded as the same location in the waggle dance, the polar coordinates were converted to Cartesian coordinates for cluster analyses. A dendrogram showing the total sum of squares for locations that the waggle dancer indicated was constructed using Ward’s minimum variance method and the statistical software R version 3.3.1. Previous research indicates that the waggle dance contains imprecise information about the vector of the food source. Weidenmuller and Seeley (1999) clarified the divergence angle in the waggle phase in relation to the distance to the trained location (θ = −0.0295 x +29.988, x: distance to the flower; θ: divergence angle in waggle phase). In this study, when the angle (Δθ) between two centroid locations of two clusters at the same distance from the hive was larger than twice the divergence angle (Δθ > 2θ), these clusters were defined as different food locations. Okada et al (2008) suggested that 80% of the waggle dance has a maximum of 30% error for the duration of the waggle phase. In this study, when concentric circles with semidiameters of 30% from two centroid locations of two clusters (that were at different distances in the same direction from the hive) overlapped, these two clusters were defined as one cluster.

Following these clustering and error corrections, the cluster locations were used to compare the locations indicated by the waggle dances of individual bees and to analyze the waggle dance developmental process.

### Statistics

We used the software R version 3.3.1 for statistical analyses. Significant differences in the duration per trip just before and just after different types of waggle dance-related behaviors were tested using the Wilcoxon signed-rank test. The trip duration among three phases after the imaginal molt was tested using the Steel-Dwass test.

## Results

### Time of appearance of waggle dance-related behaviors

How soon do waggle dance-related behaviors appear after the imaginal molt? The 100 honeybees were marked by numbered RFID tags and we calculated the number of waggle dance-related behaviors that emerged every day after the imaginal molt. Of the 100 RFID-tagged honeybees, 14 were traced until the first waggle dance. After the imaginal molt, the tagged honeybees started to go out on short round trips, less than 8 min per trip from 4 days of age (Fig. 2A). Most honeybees (n = 7) performed these short trips a few times per day; some honeybees performed the trips more than six times per day during the first week after the imaginal molt (n = 2). During this period, the honeybees performed trophallaxis (Fig. 2B) and tremble dances (Fig. 2C), but did not follow waggle dances after encountering them (Fig. 2D). One week after the imaginal molt, the tagged honeybees began to follow the waggle dance from 8 days of age (Fig. 2E and F) and performed their own waggle dance from 12 days of age (Fig. 2G and H). This suggests that dance follow precedes the waggle dance. We evaluated whether there was a common order of appearance of waggle dance-related behaviors among individuals. Waggle dance follow or one-cycle follow preceded the expression of the waggle dance in all individuals (Fig. 3A). Waggle dance follow and one-cycle follow were observed from 7 or 8 days of age and the waggle dance from 12 days of age. Individuals differed in the age at which the first waggle dance appeared (12 to 22 days of age, Fig. 3A). The interval from the first follow to the first waggle dance also differed among individuals. The shortest difference between the first follow and the first waggle dance was less than 1 day (Nos. 02 and 47); the longest difference was 8 days (No. 05).

**Fig. 2:**
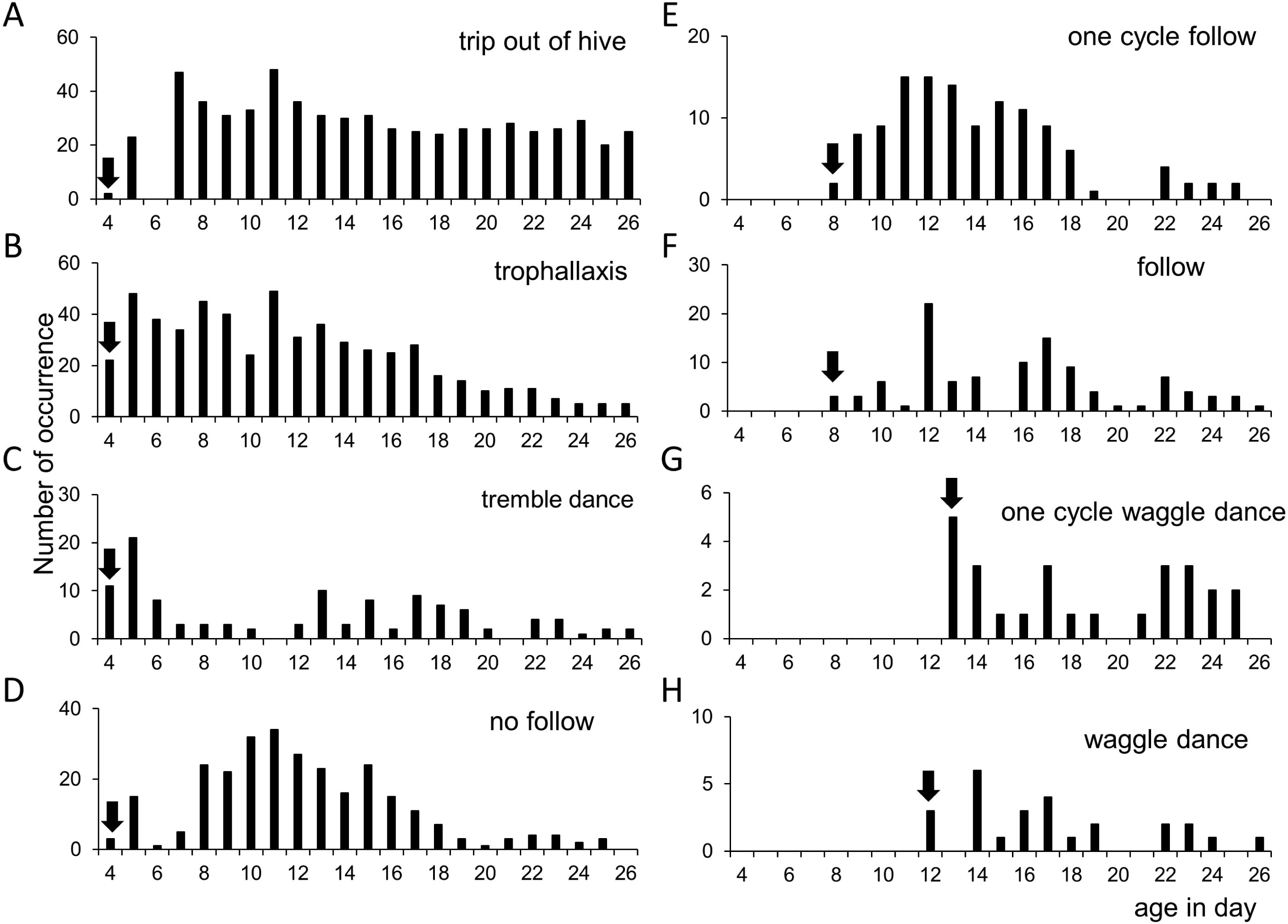
Relationship between age and numbers of each waggle dance-related behavior. Arrows indicate the day each behavior began for the most precocious individuals. From the 4^th^ day rip out of hive, trophallaxis, tremble dance were observed, however honeybees did not follow the waggle dance even if they encountered. One-cycle follow and follow were observed on the 8th day after the imaginal molt. Waggle dances and one-cycle waggle dances were observed on the 12th day and the 13th day after the imaginal molt, respectively.

**Fig. 3:**
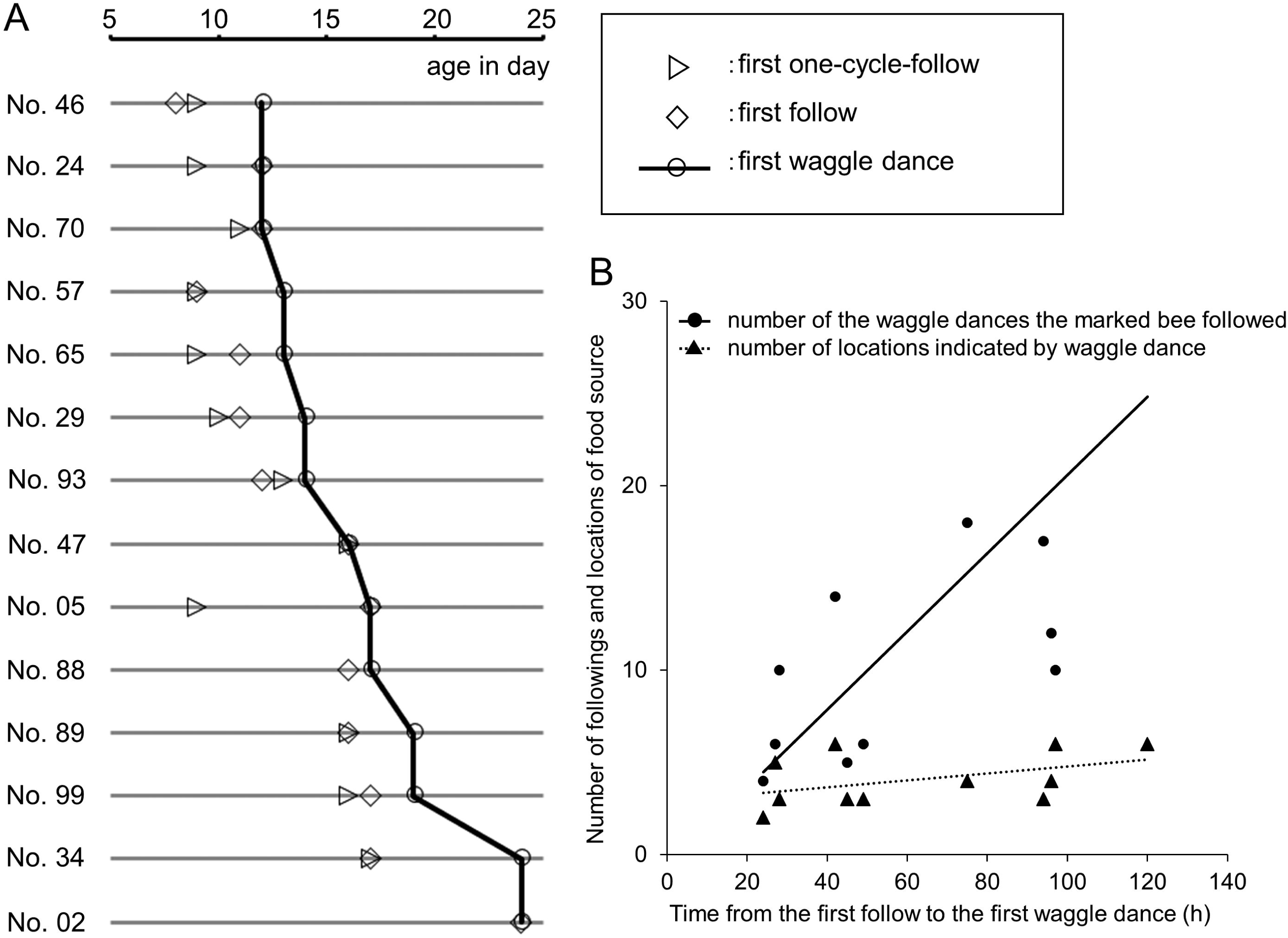
First follows and first waggle dance. A, Comparison of age (in days) of the first follow and first waggle dance for each individual (N = 14). The numbers on the left show the ID of the individual. The horizontal grey lines show the age (in days). The triangles, squares, and circles show the dates of the first appearance of one-cycle follow, follow, and waggle dances, respectively, connected by a line. In all individuals, follow preceded the first waggle dance. Individuals differed in both the age of the first waggle dance and the intervals from the first follow (or first one-cycle follow) to the first waggle dance. B, Number of follows (circles) and of food sources (triangles) in relation to the time from the first follow to the first waggle dance. The time between the first follow and the first waggle dance was positively correlated with the number of waggle dances that the marked bee followed (correlation coefficient: 0.73), but was not correlated with the number of locations indicated by waggle dances (correlation coefficient: 0.44).

What happens during the period from the first follow to the first waggle dance? We investigated the number of waggle dances each RFID-tagged bee followed and the number of locations indicated by the waggle dance during this period. The duration between the first follow and the first waggle dance was positively correlated with the number of waggle dances the bee followed (correlation coefficient: 0.73). However, there was little correlation between the duration and the number of locations indicated by waggle dances (correlation coefficient: 0.44). This suggested that the honeybees repeatedly followed waggle dances that indicated a small number of food sources (2–6 types of food source) during the period from the first follow to the first waggle dance (Fig. 3B).

Do waggle dance communications affect the following trips? Fig. 4A shows the time course of the behaviors of bee No. 29. On the 16th trip (before the first follow), the duration was less than 8 min, whereas on the 17th trip the duration substantially increased to 17 min. Subsequently, the duration was often greater than 10 min (Fig. 4A). To evaluate the effect of each waggle dance-related behavior on subsequent trips, we compared the duration per trip just before and just after the first expression of each waggle dance-related behavior. The duration per trip just before follow was significantly greater than the trip duration after follow (p < 0.01, n = 9, Wilcoxon signed-rank test); however, there were no significant differences for one-cycle follow (p = 0.125, n = 5) and waggle dances (p = 0.6257, n = 14, Fig. 4B).

**Fig. 4:**
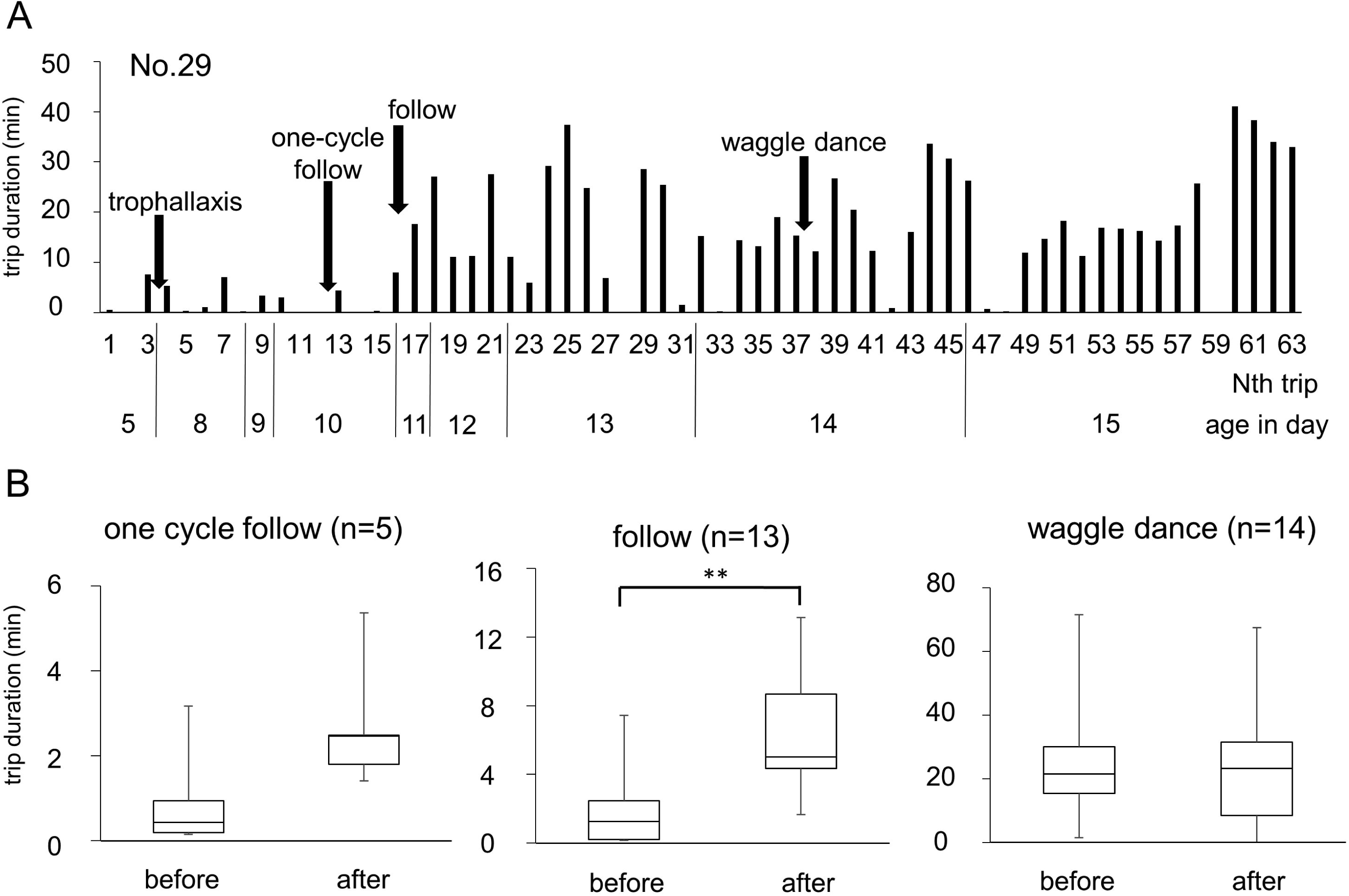
Trip duration and waggle dance-related behaviors. A, An example of the relationship between duration per trip and the first appearance of waggle dance-related behaviors of No. 29. A, The horizontal axis shows trip times on the first line and age (in days) on the second line. The vertical axis shows the trip duration. Each arrow indicates the first appearance of waggle dance-related behaviors. The trip duration substantially increased after the first follow on the 11th day of age. B, Comparison of durations of two consecutive trips just before and after each waggle dance-related behavior. The trip duration increased after the first follow (**Wilcoxon signed-rank test, p < 0.01).

### Comparison of the dance developmental process in recruits and scouts

To examine differences in the dance developmental process in recruits and scouts, we investigated the location indicated by waggle dances. To analyze the locations, we defined the vector indicated by the waggle dance of the marked honeybee as the sending vector and the vector indicated by the waggle dance that the marked honeybee followed as the received vector. If the sending vectors and the received vectors were in the same cluster, we assumed that the honeybee foraged using the information from received vectors and categorized the honeybee as a recruit (Fig. 5A). If the sending vectors and the received vectors were not in the same cluster, we assumed that the honeybee foraged without using information from received vectors and categorized the honeybee as a scout (Fig. 5B). Of 14 RFID-tagged honeybees, 11 were categorized as recruits (Nos. 02, 24, 29, 34, 57, 65, 70, 88, 89, 93, 99), and 2 as scouts (Nos. 05, 47). One of the bees (No. 46) was not categorized into these two types because although the sending and received vectors were not in the same cluster, both vectors were in the same direction (south). Therefore, it is possible that No. 46 found the location while exploring the location indicated by the received vector.

**Fig. 5:**
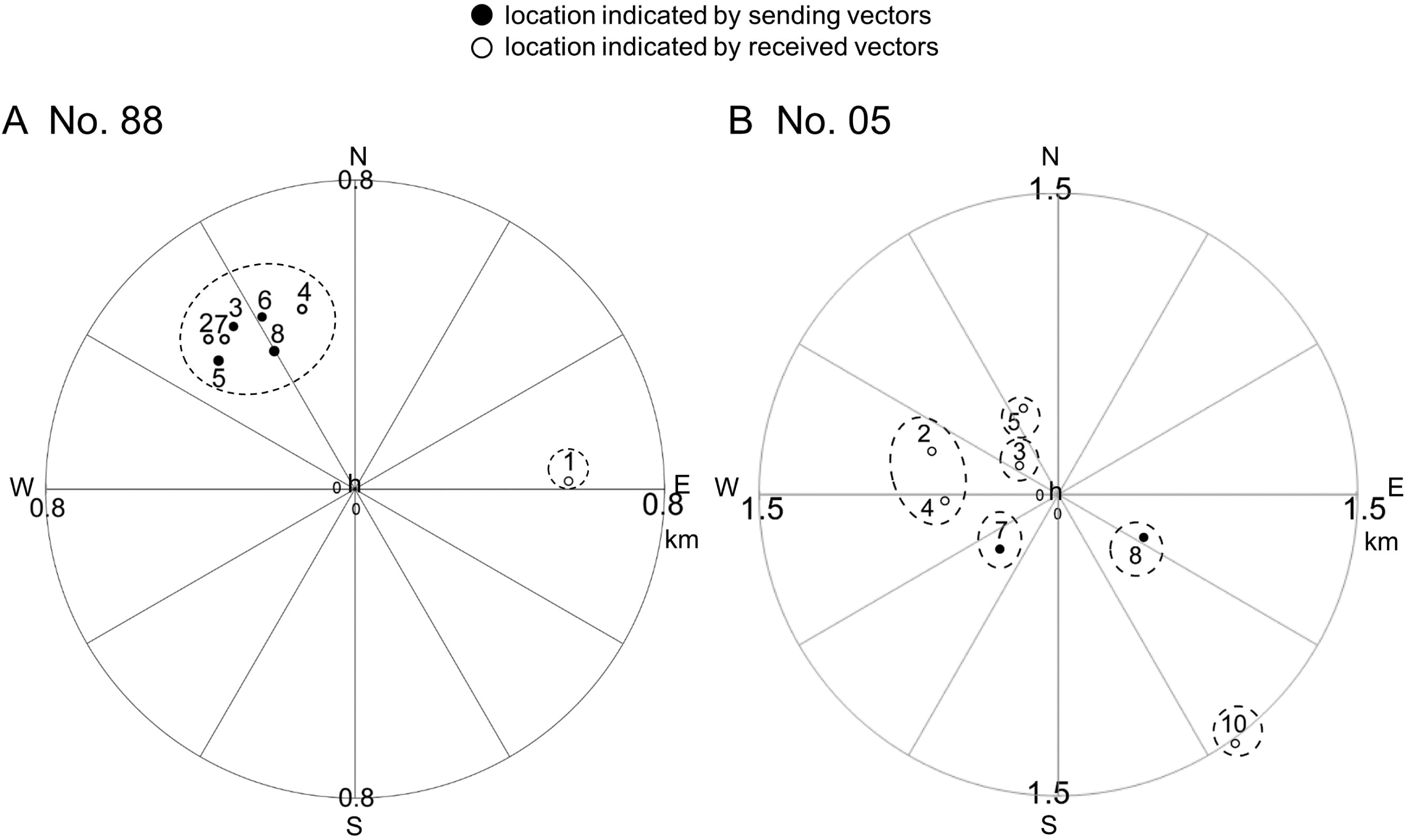
Discrimination between recruits and scouts. A, B, Map of the area around the observation hive (h; in the center). Closed dots show the locations indicated by the waggle dance of marked bees (No. 88 in A and No. 05 in B). Open dots show the locations indicated by the waggle dances that the marked bee followed previously. The numbers above the dots indicate the order in which the waggle dances appeared. The same dots which belong to the same cluster were circled by broken lines. A, The closed dots and the open dots are in the same cluster, suggesting that No. 88 was a recruit. B, The closed dots and the open dots are not in the same cluster, suggesting that No. 05 was a scout.

We investigated the duration per trip before and after the appearance of follow and the waggle dance. Fig. 6A and B show two examples of recruits and Fig. 6C one example of a scout. No. 65 twice followed waggle dances that were in the same cluster as the sending vector. After the first follow on the 9th day of age, the duration per trip was still less than 7 min. After several follows from the 11th to 12th day of age, the duration had increased remarkably to more than 20 min and on the 13th day of age the honeybee performed the first waggle dance (Fig. 6A). No. 29 followed nine waggle dances that were in the same cluster as the sending vector. The first follow was on the 10th day of age and on the 11th day of age the duration increased to more than 10 min. From the 13th day of age, the honeybee repeatedly followed waggle dances that were in the same cluster as the sending vector and flew out on trips. Finally, on the 14th day of age, the honeybee performed the first waggle dance (Fig. 6B).

**Fig. 6:**
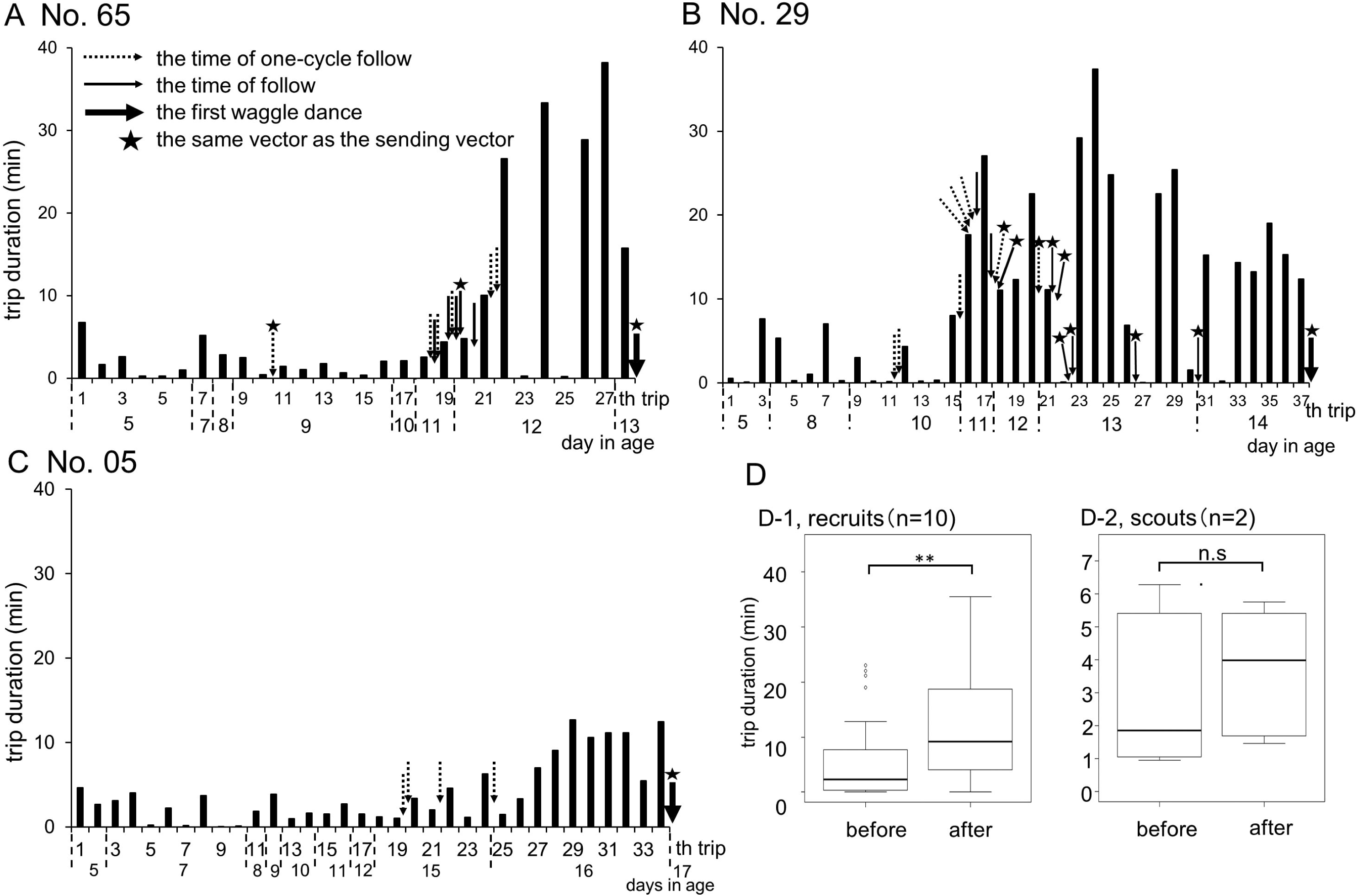
Increase in duration per trip after follow. A–C, Development of behavior before the first waggle dance in recruits (No. 65 in A and No. 29 in B) and in scouts (No. 05 in C). The horizontal axis shows the number of trips (Nth trip). The vertical axis shows the duration per trip. Dotted line arrows show times when the marked bee displayed only one follow per waggle dance. Solid arrows show times when the marked bee displayed more than two follows per waggle dance. Solid bold line arrows show the marked bee’s first waggle dance. Stars represent the same vector as that indicated by the first waggle dance. A, For No. 65, two follows indicated the same vector as that of the first waggle dance. The trip duration increased substantially from the 22nd to the 28th trip. B, For No. 29, nine waggle dance follows indicated the same vector as that of the first waggle dance. Trip durations increased substantially from the 16th to the 37th trip. C, For No. 05, the received vectors were not the same as the sending vectors. The trip duration did not substantially increase. D, In recruits, the trip duration significantly increased after the follows (D-1, Wilcoxon signed-rank test, **p < 0.01), but did not change in scouts (D-2).

For recruits, the duration per trip significantly increased after the follows (p < 0.01, n = 10, Wilcoxon signed-rank test, Fig. 6D-1). These results suggest that in recruits, waggle dance follow activates potential foraging trips and the bee uses the received vectors to explore the flowers. In contrast, the scout No. 05 made short trips of less than 5 min. On the 15th day of age, the honeybee followed only one waggle phase several times; however, the subsequent duration per trip did not increase substantially (Fig. 6C). The trip duration did not significantly increase after follows (p > 0.01, n = 2, Wilcoxon signed-rank test, Fig. 6D-2). This suggested that in scouts, waggle dance follow did not activate trips.

We divided the period from the imaginal molt to the first waggle dance of recruits (Nos. 24, 29, 57, 65, 70, 93) into Phases I, II, and III (Fig. 7A) and compared the duration per trip among these phases. There was a significant increase in the duration per trip after the first follow compared with before the first follow (Phases I vs. II, p < 0.01, n = 6, Steel-Dwass, Fig. 7B). Moreover, the duration per trip during Phase III (the period from the last follow to the first waggle dance) increased significantly more than during the other two phases (both Phases I vs. III and Phases II vs. III; p < 0.001, n = 6, Steel-Dwass, Fig. 7B). This suggested that trip duration was affected by follow and, just before the first waggle dance, affected by repeatedly following waggle dances that indicated the same received vectors.

**Fig. 7:**
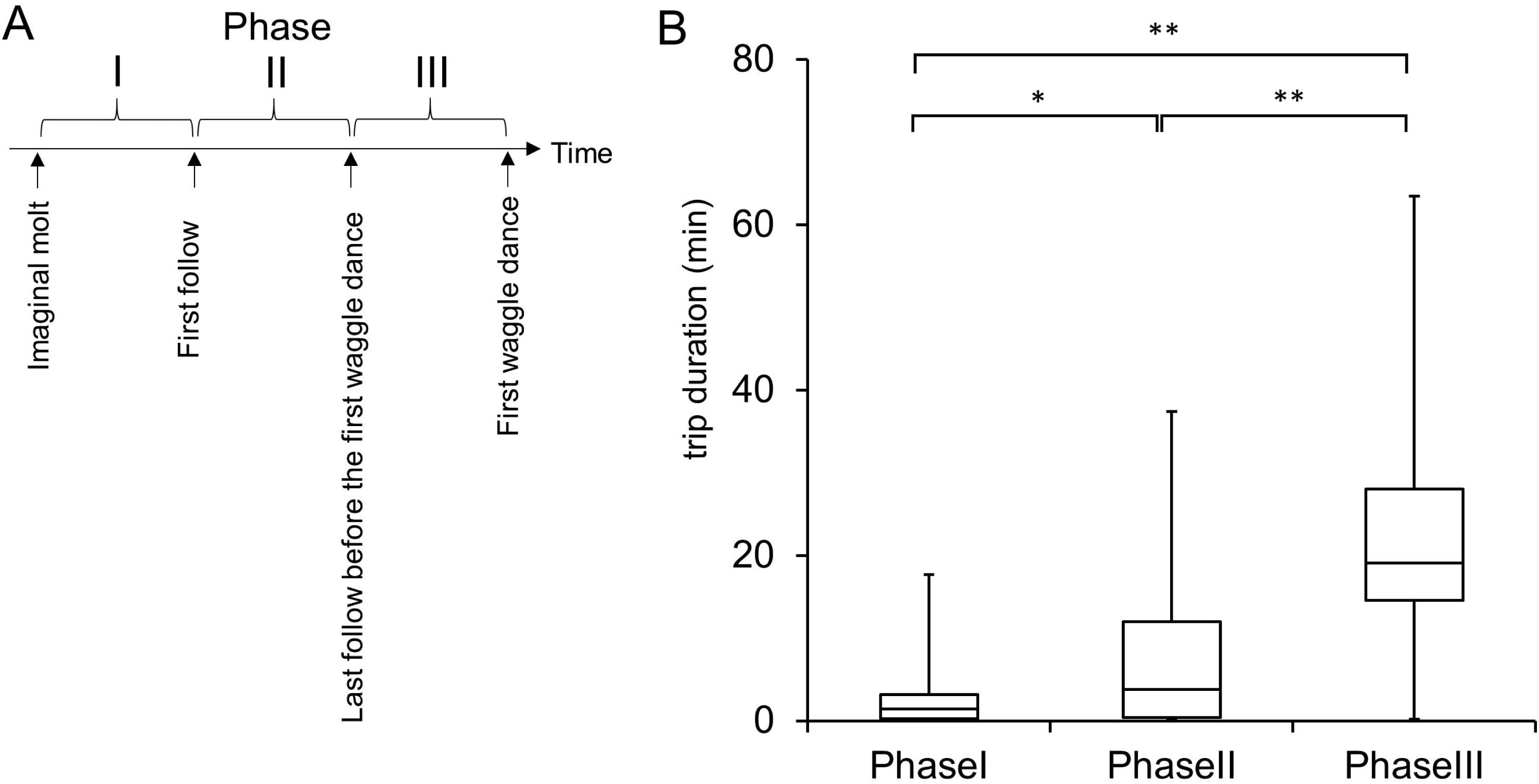
Comparison of the trip duration among the different phases before the first waggle dance for recruits. A, Three phases before the first waggle dance. Phase I is the period from the imaginal molt to the first follow. Phase II is the period from the first follow to the last follow before the first waggle dance. Phase III is the period from the last follow to the first waggle dance. B, The trip duration significantly increased in Phase III (Steel-Dwass, **p < 0.001, *p < 0.01, n = 6).

Recruits repeatedly followed waggle dances that indicated the same vector (the maximum was 12 times among 10 recruits). This suggested that recruits selectively followed waggle dances indicating a specific vector. During a waggle dance, the dancer often repeats waggle phases that indicate a similar vector. Are there any differences in the number of follows per waggle dance between a selected waggle dance and other dances? We measured the number of waggle phases that recruits followed during a waggle dance and compared them for two types of received vectors: received vectors that were in the same cluster as the sending vector (type I follow; dotted circle in Fig. 8A) and received vectors that were not in the same cluster as the sending vector (type II follow; solid circle in Fig. 8A). There were significantly more type I follow vectors than type II follow vectors (p < 0.05, n = 10, Wilcoxon signed-rank test, Fig. 8B). This suggested that recruits repeatedly followed waggle dances that indicated a specific vector and the first waggle dance indicated this vector.

**Fig. 8:**
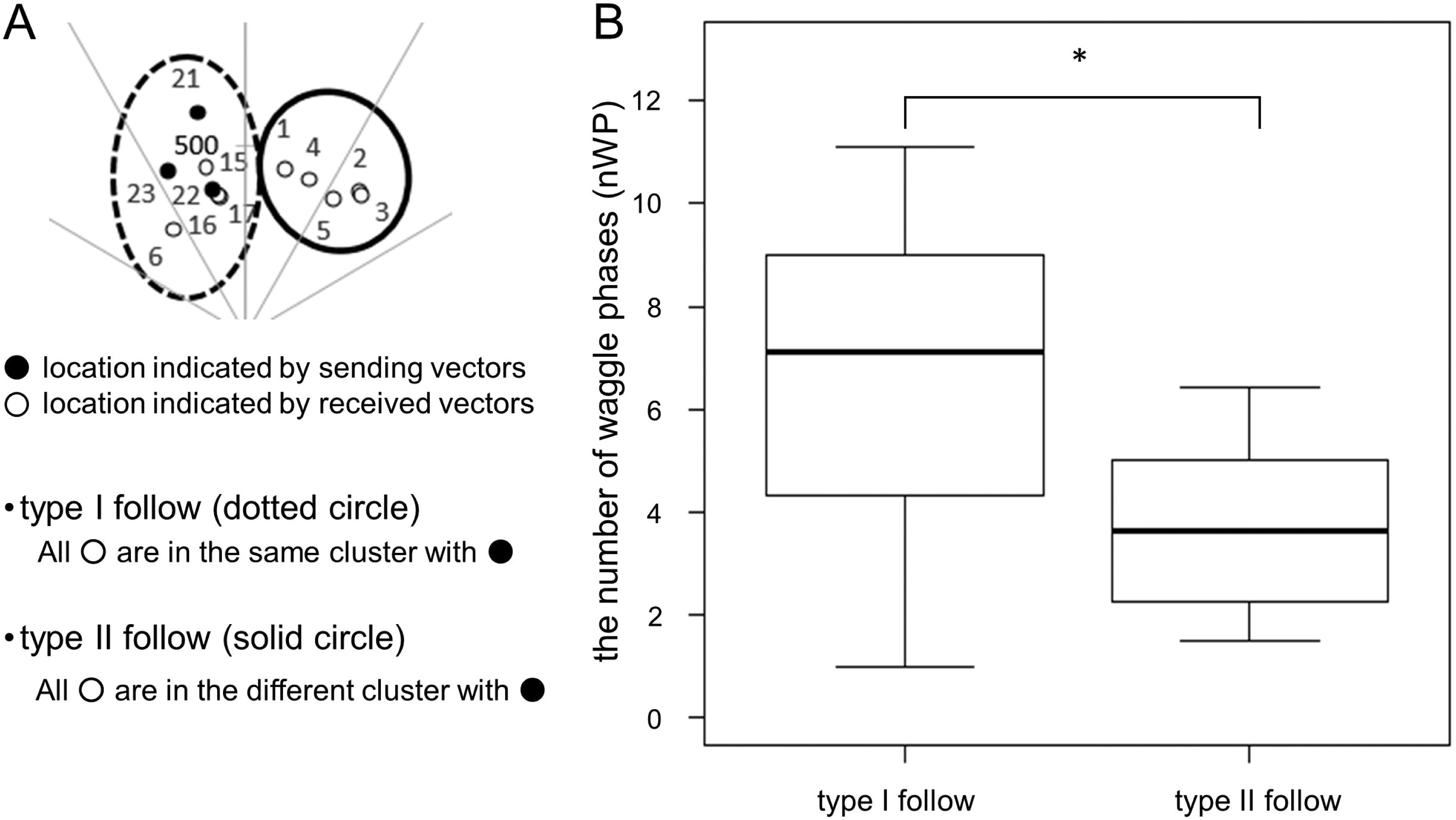
The number of follows per waggle dance for recruits. A, Two types of follows. Type I follow: The locations indicated by received vectors (open dots) are in the same cluster as those indicated by sending vectors (closed dots), which are circled by a broken line. Type II follow: The locations indicated by received vectors are in a different cluster to those indicated by sending vectors, which are circled by a solid line. B, Comparison of the number of follows per waggle dance for these two follow types. There were significantly more type I follows per waggle dance than type II follows (Wilcoxon signed-rank test, *p < 0.05, n = 10).

Can scouts convert into recruits (and vice versa)? We followed up the subsequent waggle dance history after the appearance of the first waggle dance. Of the 14 observed honeybees, 9 performed waggle dances after the first waggle dance (the other 5 bees either did not perform waggle dances or did perform waggle dances but could not be identified as recruits or scouts because the location indicated by the waggle dance was close to the other type’s cluster). We discriminated between the two types of foragers (recruits and scouts) by comparing the sending vector with previous received vectors. One of the nine bees (No. 5) was a scout on the first waggle dance (indicating location: #7, Fig. 9A). No. 5 did not subsequently follow any waggle dances and, 5 days after the first dance, performed a second waggle dance (indicating location: #8, Fig. 9A) that indicated a different cluster of received vectors. At this stage, No. 5 was still a scout.

**Fig. 9:**
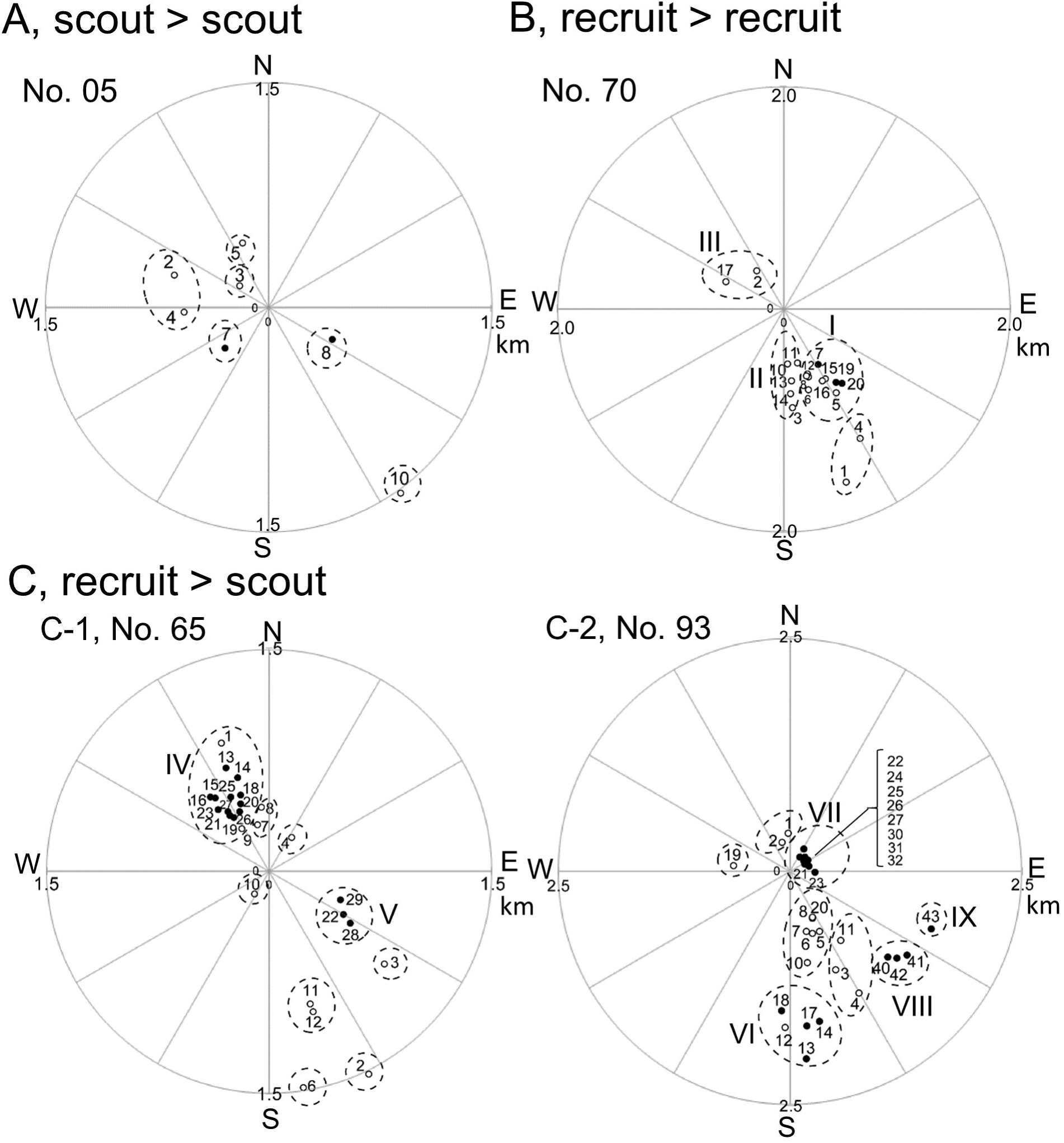
Map of the received and sending vectors of all waggle dances before and after the first waggle dance. The origin of coordinates is the location of hive. Closed dots show the location indicated by received vectors and open dots show the location indicated by sending vectors. The dots that belong to the same clusters are circled by broken lines. The number close to each dot indicates the order of appearance of the behavior (#XX). A, No. 5 was a scout on the first waggle dance (indicating location: #7) and on the second waggle dance (indicating location: #8); the dances indicated different clusters of received vectors. This bee was a scout for 5 days (see Results). B, No. 70 was a recruit on the first waggle dance (indicating location: #7 in cluster I) and the second waggle dance (indicating location: #19, #20 in cluster I), suggesting that No. 70 was a recruit for 2 days. Cluster II and III were the clusters which included the received vectors between the first and second waggle dances. C-1, No. 65 was a recruit on the first waggle dance (indicating location: #13 in cluster IV) and then changed to a scout on the second waggle dance (indicating location: #22 in cluster V). C-2, No. 93 was a recruit on the first waggle dance (indicating location: #13 in cluster VI) and then changed to a scout on the second waggle dance (indicating location: #21–32 in cluster VII). On the same day, No. 93 performed a waggle dance that indicated new locations (indicating locations: #40–43 in clusters VIII and IX). This suggests that No. 65 and No. 93 converted from recruits to scouts.

Eight of nine bees (Nos. 02, 24, 57, 65, 70, 88, 89, 93) were recruits on the first waggle dance. The location indicated by the first waggle dance of No. 70 was #7 in cluster I (Fig. 9B); subsequently, No. 70 followed waggle dances that indicated several vectors, including clusters I, II, and III (indicating locations: #8–18, Fig. 9B). The next day, No. 70 performed a waggle dance that indicated cluster I (indicating locations: #19, 20, Fig. 9B), suggesting that No. 70 was still a recruit on that day. No. 70 then disappeared, so we could not observe subsequent waggle dances.

The location indicated by the first waggle dance of No. 65 was #13 in cluster IV (No. 65 in Fig. 9C). Subsequently, No. 65 did not follow any waggle dances, made a long trip of 45 min, and repeatedly performed tremble dances after returning to the hive. After another long trip (1 hour) No. 65 performed a waggle dance that indicated another location (location #22 in cluster V of No. 65, Fig. 9C), suggesting that No. 65 explored a new location and found another food source. On the day after the first waggle dance, No. 65 made two long trips (30 min and 45 min). After returning to the hive, this honeybee showed trophallaxis and performed a waggle dance that indicated cluster IV (location #25 and #26 of No. 65, Fig. 9C); after another trip of 15 min, the bee returned to the hive and performed a waggle dance that indicated cluster V (location #22 of No. 65, Fig. 9C). This suggested that No. 65 gradually converted from a recruit to a scout.

The location indicated by the first sending vector of No. 93 was #13 in cluster VI (Fig. 9C). On the next 2 days after the first waggle dance, No. 93 followed waggle dances but did not make any trips. On the third day, No. 65 made a trip in the morning. After returning to the hive, No. 65 repeatedly performed waggle dances that indicated cluster VII (locations #21–32 in No. 93 of Fig. 9C), suggesting that No. 93 had explored a new location and found another food source. On the same day, No. 93 performed a waggle dance that indicated the new locations (clusters VIII and IX, locations #40–43 in No. 93 of Fig. 9C). This suggested that No. 93 converted from a recruit to a scout. In the following Discussion, we term this type of scout a “recruit-experienced scout” to differentiate it from a novice scout, which is a scout at the first waggle dance.

## Discussion

In this study, we clarified the waggle dance developmental process in recruits and novice scouts.

### Trophallaxis

After the imaginal molt, all observed honeybees communicated with each other most often using trophallaxis. Trophallaxis transfers olfactory information about profitable flowers to hive mates (von Frisch, 1967) and facilitates olfactory learning of hive mates (Farina et al., 2005, 2007: Gil and Marco, 2005; Farina and Wainselboim, 2005). Early olfactory experiences enhance olfactory learning and later memories of novel odors. In one study, honeybees trained at an artificial feeder with a specific odor were significantly more likely to follow waggle dances that presented the same odor 8 days later (Balbuena et al., 2012). Thus, honeybees learn food source odors through trophallaxis as early adults and then use the learned olfactory information to select waggle dances and to explore food sources. For example, we observed that No. 89 made more than six trips (maximum duration: 40 min) before the first follow. It is possible that No. 89 was exploring food sources using odor information received from previous trophallaxis. No. 89 was categorized as a recruit in the first waggle dance. Therefore, recruits might use not only vector information received from the waggle dance but also odor information for foraging.

### Waggle dance follows

The waggle dance transmits not only sending vector information but also olfactory information about a food source to hive mates (von Frisch, 1967) and dance followers use this information to explore food sources (von Frisch 1965; Mautz 1971; Bozic and Valentincic 1991; Judd, 1995). In this study, the trip duration just after follow was longer than just before follow. This indicates that the waggle dance activated the foraging of hive mates (von Frisch, 1967). Research shows that a waggle dance-mimicking robot can recruit dance followers to foraging sites (Michelsen, 1993). Therefore, the evidence suggests that waggle dance follows activate foraging, resulting in longer trips.

### Comparisons of the behavioral characteristics of recruits and novice scouts

Of the 14 bees observed in this study, 10 were categorized as recruits and 3 as novice scouts. Lindauer (1952) found that 8% of novice foragers were scouts (13/159 novice foragers), whereas 14% of novice foragers in this study (2/14) were scouts. We successfully observed the behaviors of honeybees from the imaginal molt and differentiated between novice scouts and recruit-experienced scouts. Our behavioral analyses also suggested that, on the days after the first waggle dance, some recruits performed waggle dances that indicated a new location. Lindauer (1952) also found that the population of scouts in experienced foragers was higher than in novice foragers. We found that some recruits became recruit-experienced scouts on the day after the first waggle dance (Fig. 9B).

What are the key factors that differentiate between recruits and novice scouts? To investigate this, we compared the development of the waggle dance in recruits and novice scouts. Before the first appearance of the waggle dance, recruits repeatedly followed waggle dances and then made trips of more than 10 min (Fig. 6A and B). Honeybees that have failed to forage successfully tend to follow waggle dances after returning to the hive to obtain information about food sources (Biesmeijer and Seeley, 2005; Wray et al., 2012). However, our novice scouts followed a few waggle dances and made trips of less than 10 min. The difference between recruits and novice scouts might be caused by communication ability: recruit foraging is activated by following the waggle dances. Recruits could use the information received from the waggle dance to explore food sources, whereas the foraging of novice scouts is not activated by following the waggle dance. Novice scouts cannot use the information received from the waggle dance and therefore explore food sources based on other information, such as olfactory memory from previous trophallaxis.

The recruits showed large phase-dependent changes in trip duration before the first waggle dance (Fig. 6A and B). In Phase III, the trip duration increased substantially. As the first waggle dance was performed after Phase III, this suggests the development of successful foraging activity. Moreover, recruits increased the number of follows of waggle dances that indicated the same vector as that of the first waggle dance in Phase II. The recruits received, learned, and memorized putative information about the profitable food source through repeated waggle dance follows in Phase II and explored foraging sites using this acquired information in Phase III. This suggests that recruits repeatedly confirmed the information about the flower indicated by the dancer.

### Waggle dance developmental process in recruits and scouts

In this study, we were able to trace the dance developmental process in recruits and novice scouts (Fig. 10). After the imaginal molt, honeybees make repeated short trips that include orientation flights (Capaldi et al., 2000). One week after the imaginal molt, the recruits started to follow waggle dances repeatedly and then lengthened their trip duration. They then followed waggle dances that indicated the same vector, increased the number of repeated follows per waggle dance, and lengthened the trip duration. The vector for the recruits’ first waggle dance was the same as that of the dances the recruits had repeatedly followed. In contrast, novice scouts followed the waggle dances weakly and did not increase the trip duration after dance follows. Their first waggle dance indicated a different vector to that of the dances they had previously followed. These results suggest that the difference between recruits and novice scouts may be based both on whether there is an increase in trip duration after follows. Moreover increase of the number of repeated follows to waggle dance indicated the same vector is another remarkable feature of recruits.

**Fig. 10:**
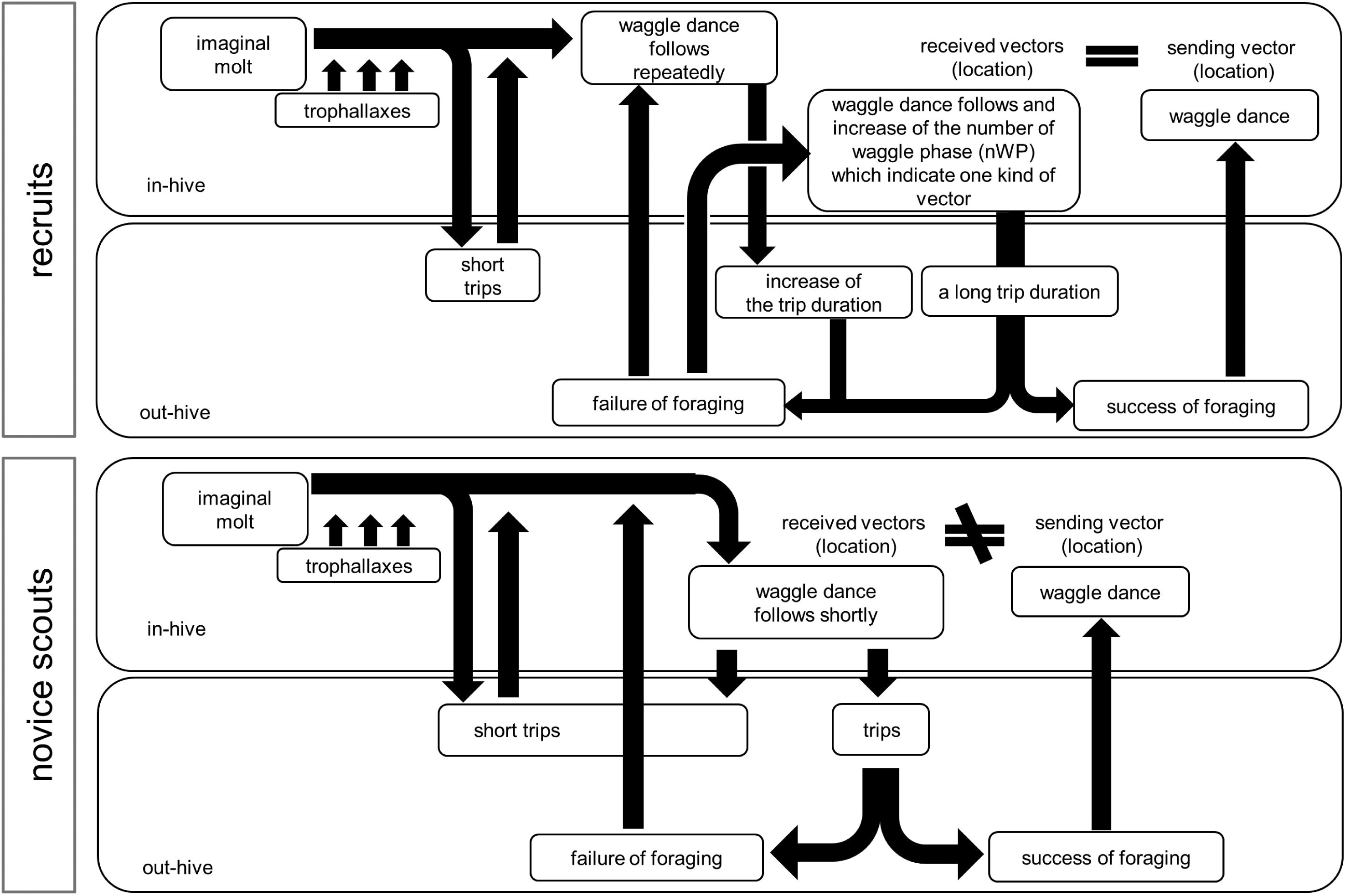
The waggle dance developmental process in recruits and novice scouts. After the imaginal molt, honeybees often show repeated trophallaxis and make short trips outside the hive (see Fig. 2). One week after the imaginal molt, recruits start to waggle dance and then increase the duration per trip (see Figs. 6A, B, and D-1). This suggests that the waggle dance activates foraging in recruits. After several trips (possibly foraging failures) recruits often repeat a specific waggle dance that indicates the same vector and engage in repeated follows (see Fig. 8). Then they make long trips and finally perform the waggle dance (see Fig. 7). The sending vector and the received vectors of the first waggle dance are the same (Figs. 6A and B). In contrast, novice scouts follow waggle dances but do not increase their trip duration (see Figs. 6C and D-2). This suggests that the waggle dance does not activate foraging in novice scouts and that the sending vector of the first waggle dance is not the same as the received vector.

In this study, we also identified recruit-experienced scouts. The developmental process of recruit-experienced scouts differs from that of novice scouts. Unlike novice scouts, recruit-experienced scouts are able to use vector information received from the waggle dance. The foragers use not only social information acquired from waggle dance communication, but also private information acquired through their own foraging experiences (Grüter et al., 2008). This exploration by recruit-experienced scouts could contribute to the discovery of new food sources. To understand the role of novice scouts in the honeybee colony, we need more detailed analyses of their characteristics, such as the location of novice scout forages, the rate of recruitment to the location, and seasonal population changes in novice scouts.

## Acknowledgments

We thank Prof Manzo Uchigasaki for technical supports for applying RFID technologies on honeybee and Prof Hidetoshi Ikeno for helpful comments on the article.

### Competing interests

No competing interests declared.

### Author contributions

All authors had full access to all study data and take responsibility for the integrity of the data and the accuracy of data analysis. Study concept and design: HA. Acquisition of data: HA, TM. Analysis and interpretation of data: HA, YK. Drafting of the manuscript: HA. Critical revision of the manuscript for important intellectual content: ST, NT. Obtained funding: HA. Setup design and programming: ST, KH, SM. Study supervision: HA.

### Funding

This work was supported by the Ministry of Education, Culture, Sports, Science and Technology of Japan, grant number 22570079 and 15K14569 and by the Central Research Institute of Fukuoka University (No. 151031 and No. 171031) to HA.

## References

Balbuena, M. S., Arenas, A., and Farina, W. M. (2012): Floral scents learned inside the honeybee hive have a long-lasting effect on recruitment. Anim. Behav, 84, 77–83.

Biesmeijer J., Seeley T. (2005): The use of waggle dance information by honey bees throughout their foraging careers. Behavioral Ecology and Sociobiology, 59, 133–142.

Boizic, J, Valentincic T. (1991): Attendants and followers of honey bee waggle dances. J Apic Res, 30, 125–131.

Capaldi, E. A., Smith, A. D., Osborne, J. L., Fahrbach, S. E., Farris, S. M., Reynolds, D. R., Edwards, A. S., Martin, A., Robinson, G. E., Poppy, G. M., Riley, J. R. (2000): Ontogeny of orientation flight in the honeybee revealed by harmonic radar. Nature, 403, 537–540.

Farina, W. M., Grüter, C., Diaz P. C. (2005): Social learning of floral odoura inside the honeybee hive. Proc Biol Sci, 272, 1923–1928.

Farina, W. M., Wainselboim, A. J. (2005): Trophallaxis within the dancing context: a behavioral and thermographic analysis in honeybees (*Apis mellifera*). Apidologie, 36, 43–47.

Farina, W. M., Grüter, C., Diaz P. C. (2007): Floral scents affect the distribution of hive bees around dancers. Behabioral Ecology and Sociobiology, 61(10), 1589–1597.

von Frisch K. (1965): Tanzsprache und Orientierung der Bienen. Springer-Verlag.

von Frisch K. (1967): The dance Language and orientation (ed. Von Frisch). Belknap Press.

Gil, M., Marco, R. J. (2005): Olfactory learning by means of trophallaxis in *Apis mellifera.* The journal of Experimental Biology, 208, 671–680.

Grüter, C., Acostona, L. E., Farina, W. M. (2006): Propagation of olfactory information withn the honeybee hive. Behav Ecol Sociobiol, 60, 707–715.

Grüter, C., Balbuena, M. S., Farina, W. M. (2008): Informational conflicts created by the waggle dance. Proceedings of the Royal Society B, 275, 1321e1327.

Judd, T. M. (1995): The waggle dance of the honey bee: which bees following a dancer successfully acquire the information? J Insect Behav, 8, 343–354.

Riley, J. R., Greggers, U., Smith, A. D., Reynolds, D. R., Menzel, R. (2005): The flight paths of honeybees recruited by the waggle dance. Nature, 435, 205–207.

Lindauer M (1952): Ein Beitrag zur Frage der Arbeitsteilung im Bienenstaat. Z Vergel Physiol, 34, 299–345.

Mautz, D (1971): Der Kommunikationseffect der Schwanzeltanzebei *Apis mellifera carnca* (Pollm.). Z Vergl Physiol, 72, 197–220.

Michelsen, A. (1993): The transfer of information in the dance language of honeybees: progress and problems. Journal of Comparative Physiology A, 173(2), 135–141.

Oettingen-Spielberg T zu (1949): Uber das Wesen der Suchbiene. Z Vergel Physiol, 31, 454–489.

Okada R, Ikeno H, Aonuma H, Ito E (2008): Biological Insights Into Robotics: Honeybee Foraging Behavior by a Waggle Dance, Advanced Robotics, 22, 1665–1681

Riley JR, Greggers U, Smith AD, Reynolds DR, Menzel R (2005): The flight paths of honeybees recruited by the waggle dance. Nature, 435, 205–207.

Seeley, T. D. (1983): Division of labor betweeen scouts and recruits in honeybee foraging. Behavioral Ecology and Sociobiology, 12, 253–259.

Seeley TD and Visscher PK (1988): Assessing the benefits of cooperation in honeybee foraging. Behavioral Ecology and Sociobiology, 22, 229–237

Thom C, Gilley DC, Hooper J, Esch HE (2007): The scent of the waggle dance. PLoS Biol 5 (9): e228.

Weidenmuller A, Seeley TD (1999): Imprecision in waggle dances of the honeybee (*Apis mellifera*) for nearby food sources: error or adaptation? Behavioral Ecology and Sociobiology, 46, 190–199

Wray, M. K., Klein, B. A., Seeley, T. D. (2012): Honey bees use social information in waggle dances foraging errors are more costly. Behavioral Ecology, 23, 125–131.

